# Quantitative proteomic profiling of tumor-associated vascular endothelial cells in colorectal cancer

**DOI:** 10.1101/561555

**Authors:** Guoqiang Wang, Qiongzhi Yang, Maoyu Li, Ye Zhang, Yu-xiang Cai, Xujun Liang, Ying Fu, Zhefeng Xiao, Minze Zhou, Zhongpeng Xie, Huichao Huang, Yahui Huang, Yongheng Chen, Qiongqiong He, Fang Peng, Zhuchu Chen

**Affiliations:** NHC Key Laboratory of Cancer Proteomics, XiangYa Hospital, Central South University, Changsha, Hunan 410008, China; Department of Pathology, XiangYa Hospital, Central South University, Changsha, Hunan 410008, China; Department of Pathology, School of Basic Medical, Central South University, Changsha 410008, China

**Keywords:** colorectal cancer, quantitative proteomics, angiogenesis, Tenascin-C

## Abstract

To investigate the global proteomic profiles of vascular endothelial cells (VECs) in the tumor microenvironment and antiangiogenic therapy for colorectal cancer (CRC), matched pairs of normal (NVECs) and tumor-associated VECs (TVECs) were purified from CRC tissues by laser capture microdissection and subjected to iTRAQ based quantitative proteomics analysis. Here, 216 differentially expressed proteins (DEPs) were identified and performed bioinformatics analysis. Interestingly, these proteins were implicated in epithelial mesenchymal transition (EMT), ECM-receptor interaction, focal adhesion, PI3K-Akt signaling pathway, angiogenesis and HIF-1 signaling pathway, which may play important roles in CRC angiogenesis. Among these DEPs, Tenascin-C (TNC) was found to upregulated in the TVECs of CRC and be correlate with CRC multistage carcinogenesis and metastasis. Furthermore, the reduction of tumor-derived TNC could attenuate human umbilical vein endothelial cell (HUVEC) proliferation, migration and tube formation through ITGB3/FAK/Akt signaling pathway. Based on the present work, we provided a large-scale proteomic profiling of VECs in CRC with quantitative information, a certain number of potential antiangiogenic targets and a novel vision in the angiogenesis bio-mechanism of CRC.

**Summery statement:** We provided large-scale proteomic profiling of vascular endothelial cells in colorectal cancer with quantitative information, a number of potential antiangiogenic targets and a novel vision in the angiogenesis bio-mechanism of CRC.

## Introduction

Tumor angiogenesis plays a vital role in creating the tumor microenvironment, and is necessary for tumor growth and metastasis. Currently, angiogenesis inhibitors have become important drugs in the treatment of solid tumors, including colorectal cancer (CRC) (Lopez *et al*, 2019; Riechelmann & Grothey, 2017). Vascular endothelial cells (VECs), which interact with tumor cells, extracellular matrix, and immune killer cells, form the major components of tumor microenvironment. Recently, antiangiogenic strategies have largely focused on targeting VECs (Liu *et al*, 2016; Liu *et al*, 2015). Tumor-associated VECs (TVECs) undergo phenotypic and epigenetic changes during tumor initiation and progression (Xiong *et al*, 2009). Increasing evidence indicates that proteins are primary targets of therapeutic drugs, and that proteins located in TVECs are potential therapeutic targets against tumor angiogenesis and widely used in screening antiangiogenic drugs (Kalen *et al*, 2009; Sonveaux, 2008).

Proteomics has introduced an effective cancer research method that provides new opportunities for discovering the therapeutic targets of CRC and for revealing the underlying molecular mechanism of this disease. The proteomics of CRC has been extensively studied (Alvarez-Chaver *et al*, 2018; Peng *et al*, 2016), but relatively few studies have focused on tumor microenvironment, especially the proteome of VECs in CRC. However, the VECs comprises only a relatively small percentage of the total tumor volume. Consequently, an altered VECs expression signature may be easily masked in whole tumor studies. To overcome this limitation, laser capture microdissection (LCM) was exploited to isolate a relatively pure VECs from heterogeneous frozen tumor tissue samples in this study.

We compared protein expression in VECs isolated by LCM from CRC tumors and matched adjacent nonmalignant colorectal (ANC) tissues from the same patient. This approach has the advantage of eliminating some of the inherent heterogeneity between individual patients and between different cell types present in the samples (Johann *et al*, 2010; Unwin *et al*, 2003). The resulting peptides of the enzymatically digested proteins were measured by iTRAQ-based quantitative proteomics. Compared to conventional proteomic technology, this approach possesses many advantages, such as high throughput, high accuracy, high repeatability, and high sensitivity, and it can be used for various types of biological samples (Pierce *et al*, 2008). Based on our data, we presented the proteome of VECs in CRC using LCM technology and quantitative proteomics analysis. These TVECs-related proteins may serve as potential therapeutic targets and increase our understanding of the mechanisms underlying CRC angiogenesis.

## Results

### Proteomic profiling of VECs in CRC

We compared the global proteomic profiles of paired TVECs and NVECs in 10 patients with CRC to identify proteins and pathways that may be associated with the CRC angiogenesis. As a result, A total of 2058 nonredundant proteins were repeatedly identified and quantified at a minimum confidence level of 95% (unused ProtScore > 1.3) by triplicate iTRAQ labeling and 2D LC-MS/MS analyses (Supplementary Table S1). A protein density plot was subsequently generated to determine the thresholds for clustering DEPs using the ratios of those quantified proteins (Chakraborty *et al*, 2015). Using 10%, 90% and in-between quantile-based thresholds, averaged ratio-fold changes >1.2129 or < 0.7963 between two cohorts of proteins were categorized as upregulated and downregulated proteins, respectively (Figure 1A).

**Figure 1:**
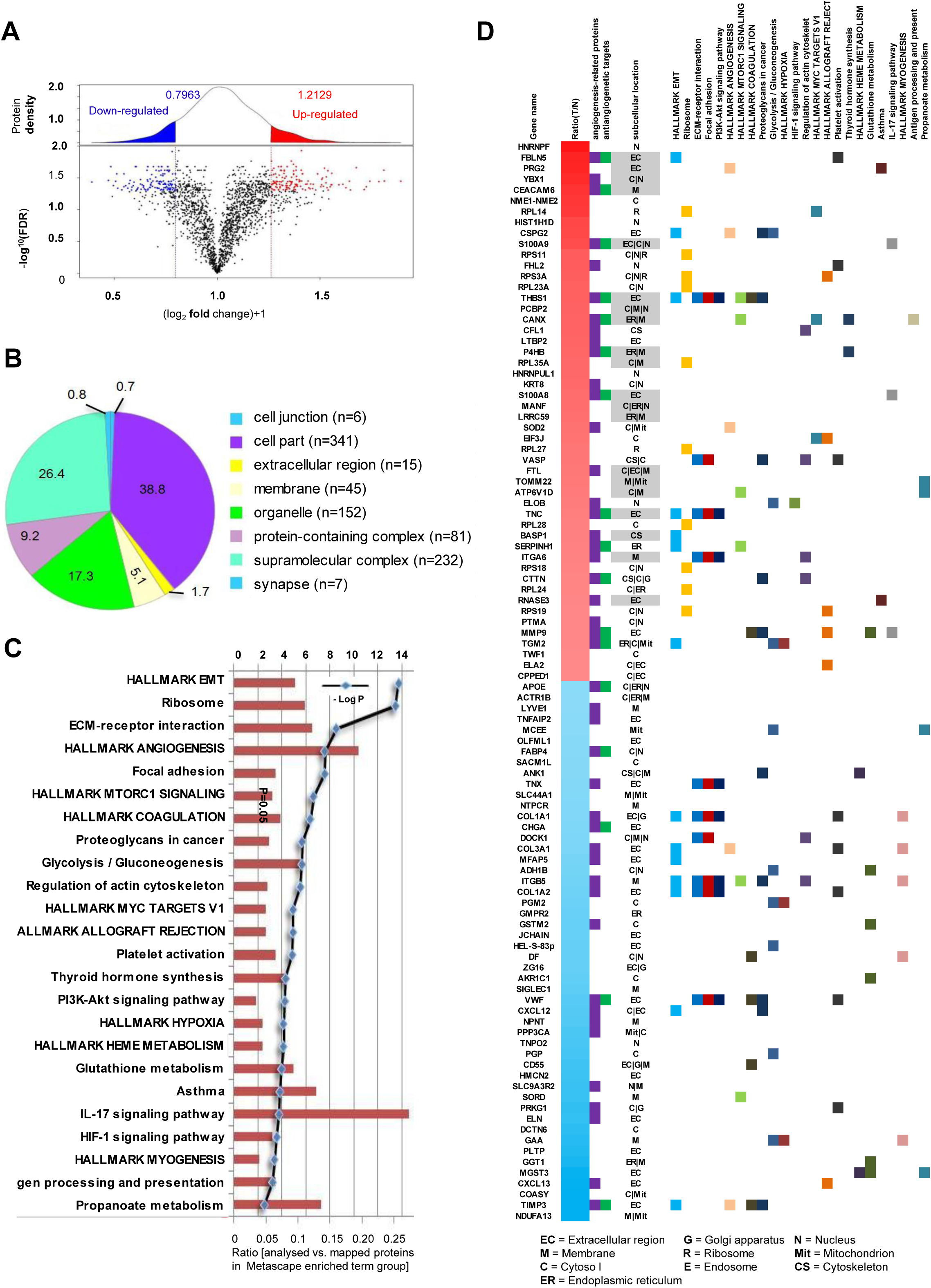
Integrative analysis of identified proteins. (A) The ratio intensity plot representing protein fold change (iTRAQ ratio versus corresponding summed peptide intensity distribution) and protein density plot (upper panel). Red, blue and black clusters indicate up-, down- and unregulated proteins, respectively. (B) A total of 2058 proteins was classified according to the cell components with PANTHER. (C) Metascape pathway analysis mapped the DEPs to 24 signaling pathways. (D) Heatmap of top 50 up- and top 50 down-regulated proteins mapping to each Metascape pathway terms group. The subcellular location of each protein is also presented and up-regulated proteins that are either secreted or membrane are highlighted in gray as potential therapeutic targets.

A total 216 proteins were found to be differentially expressed, including 119 upregulated proteins and 97 downregulated proteins in TVECs relative to NVECs (Supplementary Table S2). Compared to previous literatures, 45 of the top 100 DEPs (top 50 up- and top 50 down-regulated proteins) have been proved as angiogenesis-related proteins, and 17 of the 45 proteins have been reported or predicted as potential targets for cancer antiangiogenic treatment. These proteins are presented in heatmap format in (Figure 1D). In addition, the top 50 up-regulated proteins that are either secreted or localised in the membrane are highlighted in gray in the heatmap as potential therapeutic targets in TVECs (gene names of the respective proteins are: FBLN5, PRG2, YBX1, CEACAM6, S100A9, THBS1, PCBP2, CANX, P4HB, RPL35A, S100A8, MANF, LRRC59, FTL, TOMM22, ATP6V1D, TNC, BASP1 and RNASE3) (Figure 1D). To some degree, these results indicate that our findings were consistent with previous studies and also make new discoveries.

### Bioinformatics Analysis

To obtain a biological view of the identified proteins, a total of 2058 proteins were classified according to cellular compartment levels using the PANTHER GO classification system. Variable cellular compartments cell part (38.8%), organelle (26.4%) and membrane (5.1%) are showed in (Figure 1B). Metascape enrichment analysis, accounting for all the DEPs, demonstrated that processes related EMT, ECM-receptor interaction, focal adhesion, PI3K-Akt signaling pathway, angiogenesis and HIF-1 signaling pathway were over-represented (Figure 1C and Supplementary Table S3). The top 50 upregulated and downregulated proteins mapping to each Metascape enrichment term group are presented in heatmap format in Figure 1D. The subcellular localization of these proteins is also presented in the heatmap. Moreover, Metascape enrichment analysis mapped all the identified proteins to 66 signaling pathways (Supplementary Table S4). Interestingly, the results demonstrated that TVECs exhibited an increased dependence on processes related with focal adhesion, PI3K-Akt signaling pathway, HIF-1 signaling pathway and EMT (Figure 2A, 2B and Supplementary Figure S1, Table S5), which might play an important role in CRC angiogenesis.

**Figure 2:**
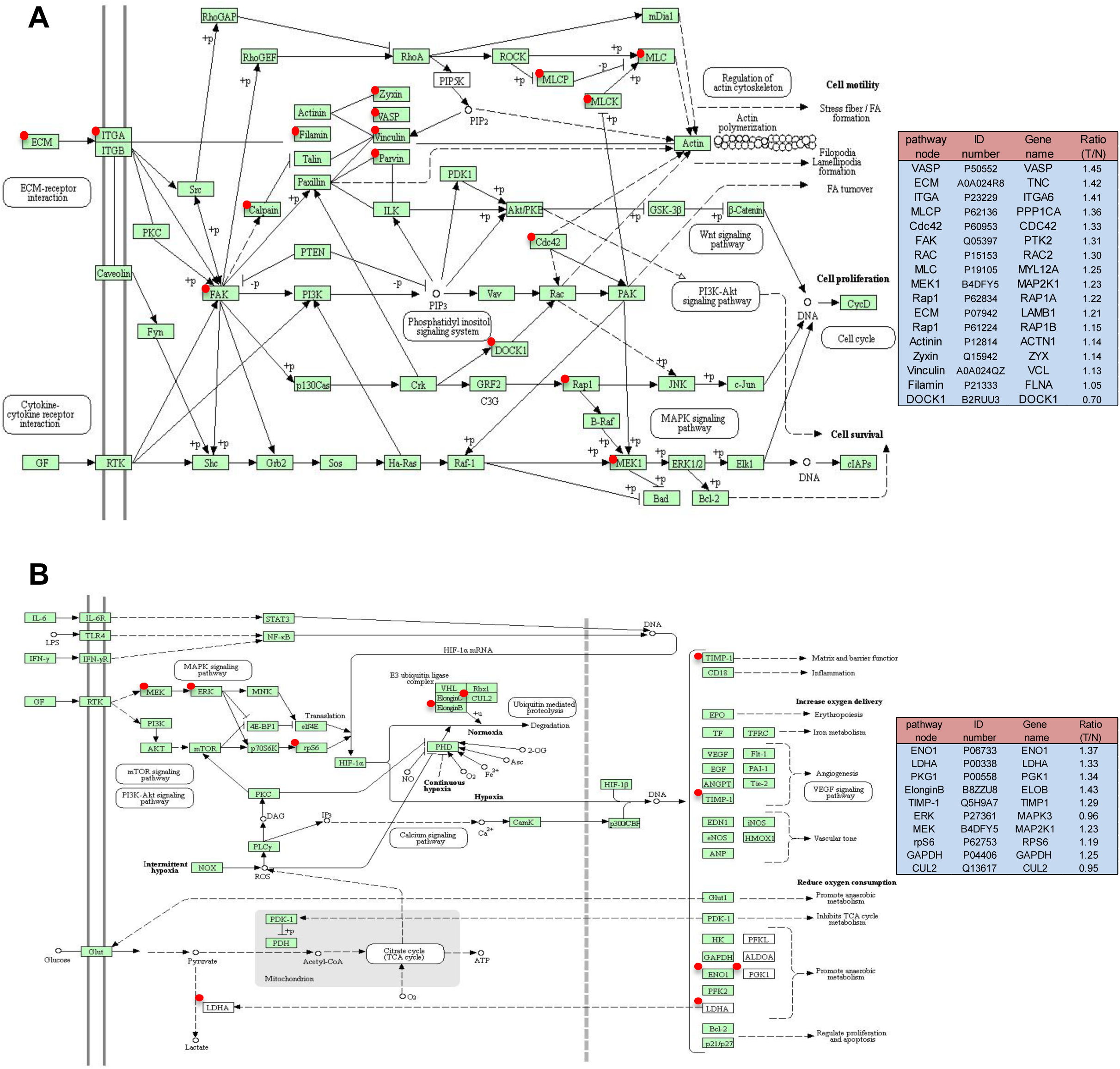
Focal adhesion pathway (A) and HIF-1 signaling pathway (B) altered in a CRC. Green rectangle with red mark means the identified proteins. Green rectangle without red mark means species-specific enzymes. White rectangle means reference pathway. The solid line indicates molecular interaction. The dot line means indirect effect. The pathway node in the right panel corresponds to the red marked node in the left diagram. ID number is the Swiss-Prot accession number. Ratio (T/N) = Ratio of TVECs to controls.

### Validation of the expression of TNC in VECs using immunohistochemistry

In our study, we found TNC was involved in focal adhesion, PI3K/Akt signaling pathway and EMT. To confirm the expression and location of TNC in CRC tissues, we detected the expression of TNC using immunohistochemistry in 30 cases of NCM, 30 cases of AD, 30 cases of CIS and 50 cases of ICC. As shown in Figure 3 and Table 1, the expression levels of TNC were progressively increased in the CRC carcinogenic process from early stage, AD, to late stage, ICC. (P<0.05). Strong TNC immunostaining was readily detected in the stoma and VECs of the CIS and ICC (Figure 3A and 3B), whereas weak staining in AD and negative staining in NCM were detected (Figure 3C and 3D). In addition, the expression levels of TNC in VECs of lymph nodes with metastasis were strong compared to those in lymph nodes without metastasis (Figure 3E and 3F). Immunostaining of NCM and ICC for the EC marker CD34 was used as a positive control for VECs (Figure 3G and 3H).

**Figure 3:**
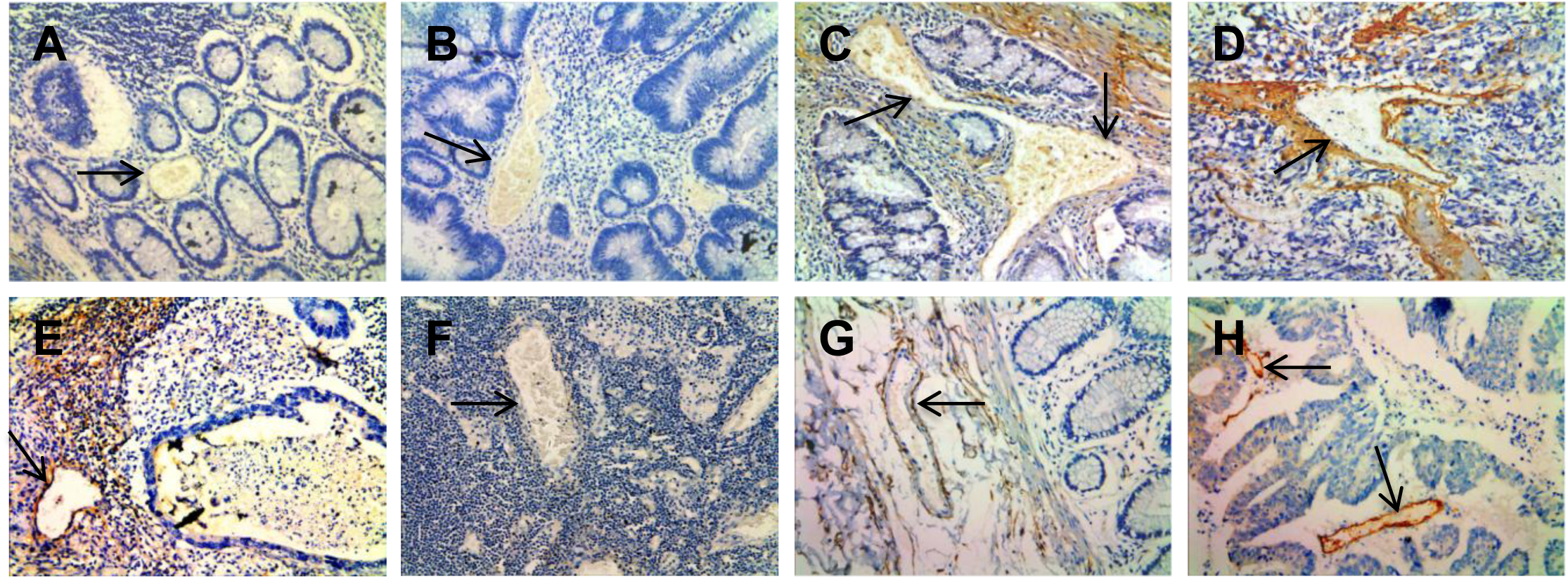
Representative results of immunohistochemistry show the expression of TNC in vessels. Original magnification, × 200. Top panel: TNC immunostaining of NCM (A), ACP (B), CIS (C) and ICC (D). Negative staining was observed in VECs of NCM and ACP, moderate staining in CIS, and strong staining in ICC. Bottom panel: Strong staining was observed in VECs of lymph nodes with metastasis (E). Negative staining was observed in VECs of lymph nodes without metastasis (F). Endothelial cell marker (anti-CD34) immunostaining of NCM (G) and ICC (H).

**Table 1.** TNC expression in multistage of colorectal epithelial carcinogenic vessels

| Multistage | Number | TNC |  |  | P-value |
| --- | --- | --- | --- | --- | --- |
|  |  | Low ( 0-2 ) | Medium( 3-4 ) | High ( 5-6 ) |  |
| NCM | 30 | 23 | 6 | 1 | P < 0.0001 |
| AD | 30 | 15 | 13 | 2 |  |
| CIS | 30 | 4 | 17 | 9 |  |
| ICC | 50 | 7 | 16 | 27 |  |

Notes: P<0.05 by linear trend chi square test, NCM vs. AD vs. CIS vs. IC.

Furthermore, we examined the relationship between the expression levels of TNC and clinicopathological characteristics in the 80 cases of CRC tissues above (30 cases of CIS and 50 cases of ICC). The results showed that TNC expression levels in VECs of CRC were closely correlated with lymph nodes metastasis (P < 0.05) and distant metastasis (P < 0.01) but had no relation with age or gender (P > 0.05; Table 2).

**Table 2.**
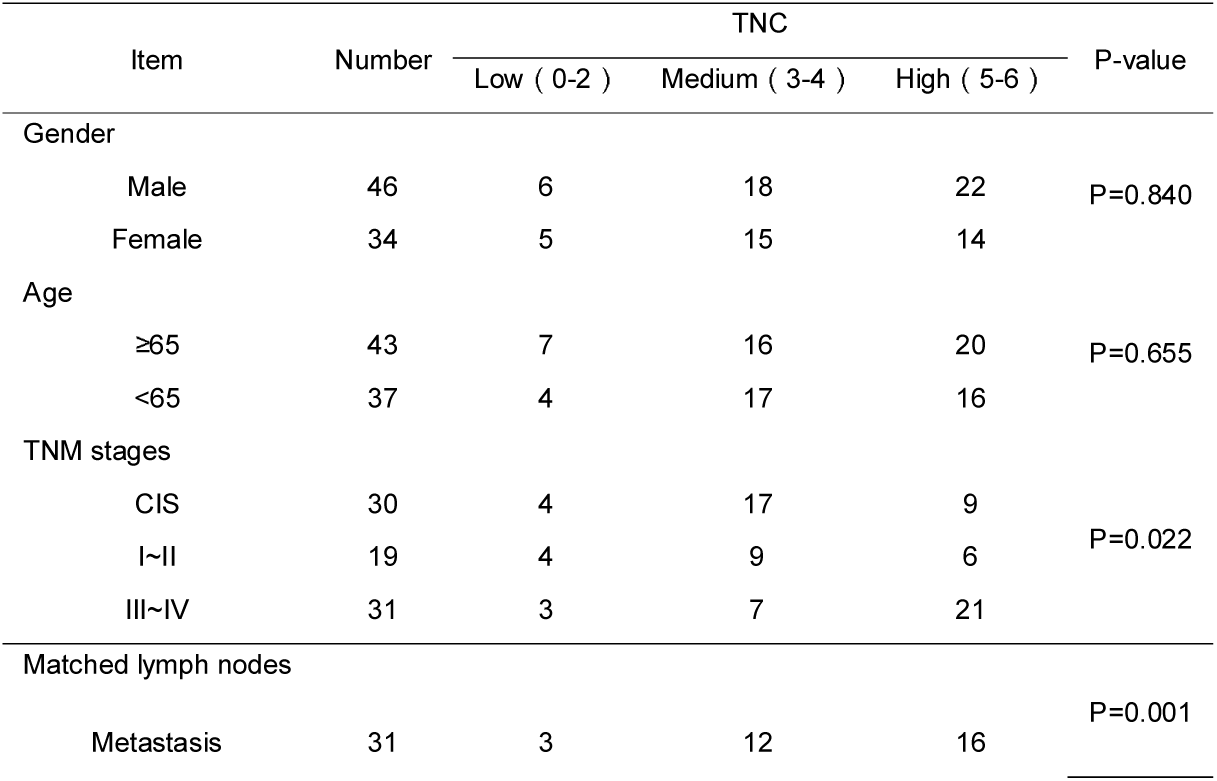
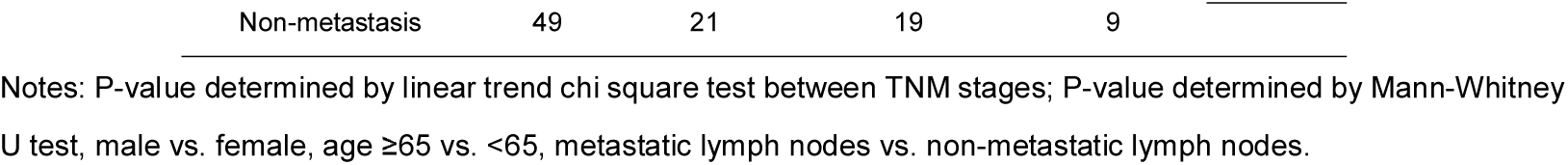
Correlation between expression of TNC protein and clinicopathological characteristics of patients with colorectal carcinoma

### Reduction of tumor-derived TNC inactivate ITGB3/FAK/Akt signaling

Tumor-derived TNC promotes or enhances the process of angiogenesis in different tumor models (Hirata *et al*, 2009; Kawamura *et al*, 2018; Pezzolo *et al*, 2011; Rupp *et al*, 2016), but the mechanistic insight has not been fully elucidated. In our previous study, TNC level is much higher in high metastatic potential CRC cell line SW620 compared to the other four CRC cell lines, particularly in the conditioned medium (Li *et al*, 2016). In the current study, We generated CRC cell line SW620 with knockdown of TNC (Figure 4A and 4B), then collected conditioned media from SW620, SW620/Vector and SW620/shTNC cells and subsequently cultured HUVECs with the conditioned media for 24 h. The results indicated that conditioned media from SW620/shTNC cells reduced the expression of ITGB3, Phospho-FAK Tyr397, Phospho-Akt Ser473 (Figure 4C), however, it did not alter the expression of TNC (Figure 4C). These results demonstrated that tumor-derived TNC has a positive influence on ITGB3/FAK/Akt signaling pathway.

**Figure 4:**
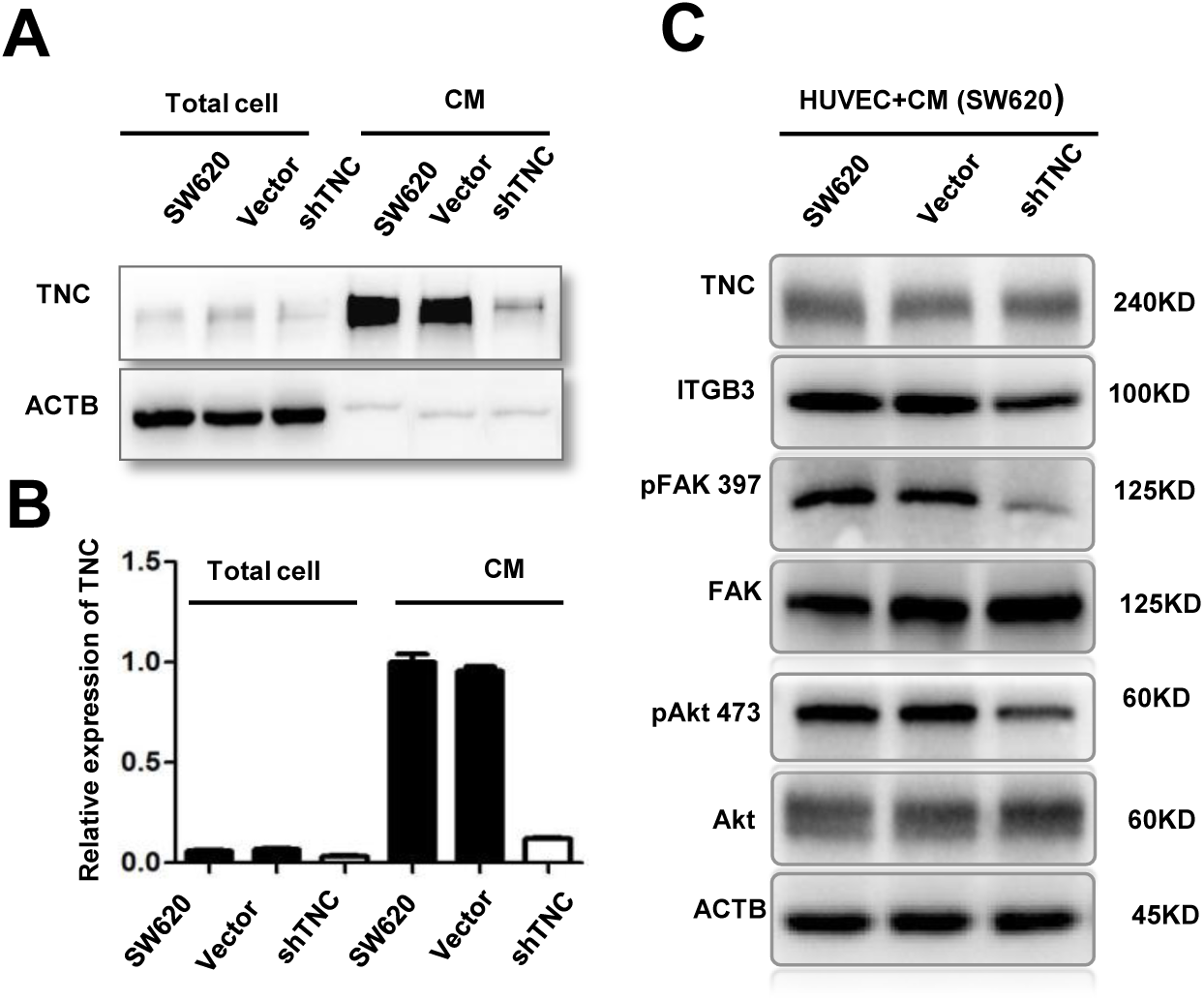
Reduction of tumor-derived TNC inactivate ITGB3/FAK/Akt signaling in HUVECs. (A-B) TNC protein levels in total cell and conditioned medium from untransfected (SW620), empty vector (SW620/Vector) and TNC-shRNA plasmid transfected SW620 cells (SW620/shTNC) were detected by western blot analysis. (C) Treatment with conditioned media (CM) from SW620/sh-TNC cells decreased ITGB3, phosphorylation of FAK-397 and phosphorylation of Akt-473 in HUVECs as compared with CM from untransfected SW620 cells.

### Reduction of tumor-derived TNC impairs tubulogenesis, proliferation and migration of HUVECs in vitro

To further confirm whether TNC has an impact on tubulogenesis activity of HUVECs, we plated HUVECs on matrigel together with conditioned media from SW620, SW620/Vector and SW620/shTNC cells respectively. As shown in Figure 5A and 5B, when TNC knockdown, the incubation of HUVECs with SW620/shTNC conditioned media resulted in a 60% decrease in the formation of capillary-like structures compared with the tubules formed by HUVECs incubated with SW620 conditioned media, and similar results were obtained in cell proliferation and migration of HUVECs using CCK8, wound-healing and transwell chamber assays (Figure 5C, 5D, 5E, 5F and 5G). Our results suggested that TNC has a positive influence on the proliferation, migration and tubulogenesis of HUVECs.

**Figure 5:**
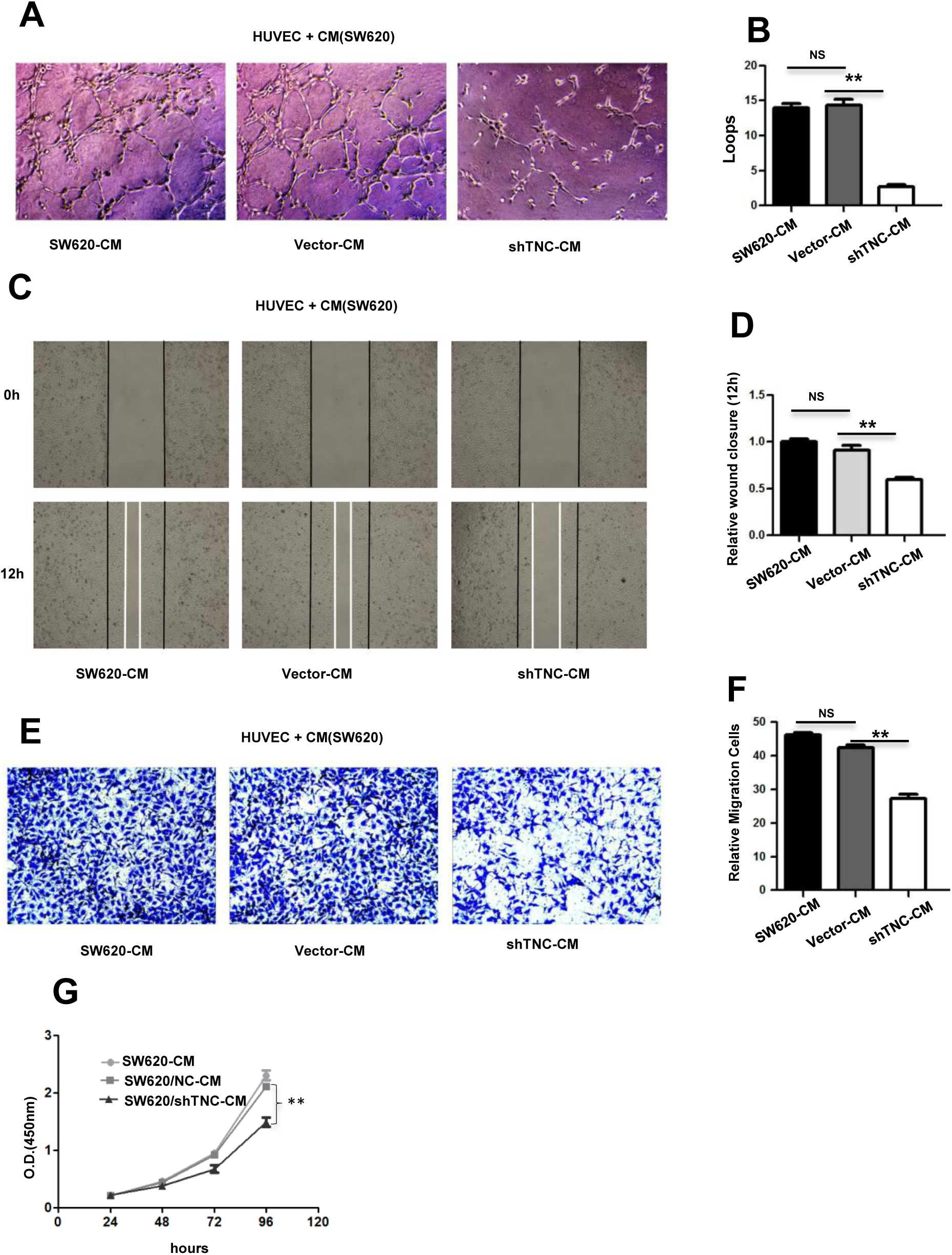
Reduction of SW620-derived TNC leads to the decease in proliferation, tubulogenesis and migration of HUVECs *in vitro* (A-B) Representative images (A) of HUVECs 7 h after plating on Matrigel together with conditioned medium from untransfected (SW620-CM) cells and empty vector (Vector-CM)- and TNC-shRNA plasmid-transfected SW620 cells (shTNC-CM) followed by quantification of the number of endothelial closed loops (B). Values are mean ± SEM from three independent experiments with three replicates. (C-D) Wound closure of HUVECs at 12 h was quantified upon addition of SW620-CM, Vector-CM and shTNC-CM. Values are the mean ± SEM from three independent experiments with three replicates. (E-F) Representative pictures (E) and quantification (F) of HUVEC migration through a transwell chamber containing SW620-CM, Vector-CM and shTNC-CM after 24 h. Values are the mean ± SEM from three independent experiments with three replicates. (G) CCK8 assay of HUVECs after treatment with SW620-CM, Vector-CM and shTNC-CM up to 96 h. Values are the mean ±SEM in HUVECs from three independent experiments with five replicates.

## Discussion

Tumor angiogenesis is a complex process leading to abnormalities in vascular structure and function (Sasaki *et al*, 1991). Similar to vasculature in individual organs, TVECs are tissue-specific, which is mostly depending on the tumor microenvironment (Jin *et al*, 2016). In this study, we compared global proteome profiles of VECs derived from normal and tumor tissues to gain insight into CRC angiogenesis and discover antiangiogenic targets for tumor therapy. A total of 216 DEPs were identified, and then GO analysis and systematic pathway-based enrichment analysis was performed. Among the DEPs, many have been reported or predicted as potential targets for cancer antiangiogenic treatment (see Figure 1D and Supplementary Table S4), such as FBLN5 (Albig & Schiemann, 2004), CEACAM6 (Zang *et al*, 2015), S100A9 (Eisenblaetter *et al*, 2017; Zhang *et al*, 2017), THBS1 (Lawler, 2002), CANX (Demeure *et al*, 2016), TNC (Kawamura *et al*, 2018), HSP47 (Wu *et al*, 2016), CTTN (Ramos-Garcia & Gonzalez-Moles, 2018), MMP9 (Gupta *et al*, 2013), TGM2 (Lei *et al*, 2018), S100A7 (Padilla *et al*, 2017), LCN2 (Hu *et al*, 2018), RACK1 (Wang *et al*, 2011), PGK1 (Shichijo *et al*, 2004), EPO (Samoszuk *et al*, 1996), CD74 (Gai *et al*, 2018), GRP78 (Kao *et al*, 2018). For these candidate antiangiogenic targets, our results are consistent with previously published data. Furthermore, the focal adhesion, PI3K-Akt signaling pathway, HIF-1 signaling pathway and EMT were identified as significantly and consistently proangiogenic categories in CRC as these pathway-related proteins were significantly upregulated in TVECs compared to controls.

Focal adhesion is a subcellular structure which acts as a scaffold for many signaling pathways involving integrin or the mechanical force exerted on cells (Shen *et al*, 2018). Many molecules in the focal adhesion complex are implicated in downstream signaling pathways, such as the AKT1 (Higuchi *et al*, 2013), MAPK/ERK pathway (Ye *et al*, 2017), and Wnt signaling (Yu *et al*, 2012). In this way, pathways impacted by the focal adhesion complex are as varied as apoptosis (Bouchard *et al*, 2008), cell proliferation (Luo *et al*, 2018), cell migration and angiogenesis (Zhao & Guan, 2011). In our findings, the focal adhesion-related proteins were significantly increased in TVECs as compared to controls (Figure 2A), which indicated the focal adhesion pathway may play a crucial role in CRC angiogenesis. Therefore, uncovering the molecular processes underlying focal adhesion hub signaling will foster the development of reasonable and feasible multimodal treatment options towards CRC.

The role of PI3K/Akt signaling pathway and HIF-1 signaling pathway in angiogenesis and tumor progression were well-documented (Karar & Maity, 2011). Hypoxia leads to the stabilization of HIF-1α and is a major stimulus for tumor cells to increase the expression of VEGF. However, the activation of the PI3K/AKT pathway in tumor cells can also increase VEGF secretion. Moreover, PTEN/PI3K/AKT regulates the proteasome-dependent stability of HIF-1α under hypoxic conditions and controls tumor-induced angiogenesis and metastasis (Joshi *et al*, 2014). The HIF-1signaling pathway and PI3K signaling pathway have been exploited for the development of new cancer therapies (Post *et al*, 2004; Tanaka *et al*, 2015; Thorpe *et al*, 2015). In the current study, HIF-1 and PI3K/AKT signaling pathway-related proteins were significantly upregulated in TVECs compared with controls (Figure 2B and Supplementary Figure S1). Those findings indicate that PI3K and HIF-1 inhibitors are excellent candidates for the treatment of CRC.

EMT underlies the progression and metastasis of malignant tumors by enabling cancer cells to depart from their primary tumors, invade surrounding tissues and spread to distant organs. Endothelial-to-mesenchymal transition (EndMT) is often categorized as a specialized form of EMT. Recent studies suggested that EndMT may play a role in angiogenic sprouting by enabling the so-called tip cells, which lead an emerging vascular plexus, to migrate into adjacent tissue (Potenta *et al*, 2008; Welch-Reardon *et al*, 2015). In our study, EMT-related proteins were significantly increased in TVECs as compared to controls (Supplementary Table S5). For example, FBLN5 (fold change=1.81) initiates EMT and enhances EMT induced by TGF-beta in mammary epithelial cells via a MMP-dependent mechanism (Lee *et al*, 2008). Based on the clinical application of anti-angiogenesis therapy in metastatic CRC, the EMT-angiogenesis and EMT stemness links in the CRC cells, newly synthesized drugs with antiangiogenic/anti-EMT properties could be one of the medications used in the future in the targeted therapy of patients with CRC (Gurzu *et al*, 2016).

In our findings, TNC was implicated in focal adhesion, PI3K/Akt signaling pathway and EMT. Several researchers have demonstrated that TNC promotes neoplastic angiogenesis in various cancers including breast cancer, glioblastoma, and oral and pharyngeal squamous cell carcinoma (Atula *et al*, 2003; Mai *et al*, 2002; Rupp *et al*, 2016). However, few reports have investigated the expression of TNC in VECs of CRC and the mechanism for promoting CRC angiogenesis. In our study, we found that the expression of TNC in VECs correlated with CRC multistage carcinogenesis. This finding suggested that TNC may have a positive effect on CRC angiogenesis and progression. As a following step, we highlighted the role of TNC in triggering ITGB3/FAK/Akt-473 signaling pathway, which promoted HUVEC migration and angiogenesis. Moreover, TNC can be secreted in large amounts by SW620 cells (Li *et al*, 2016), indicating that TNC is more likely to end up in the blood or other body fluids in a ‘measurable’ concentration in CRC patients, which can be used for blood-based diagnostics. Therefore, we demonstrated the potential regulation mechanism of TNC and indicated its possible use as a diagnostic and therapeutic target in CRC.

In conclusion, our study built the first proteome of VECs in CRC and regarded the role of vascular proteins in the process of colorectal angiogenesis. In addition, we demonstrated the function of TNC in regulating ITGB3/FAK/Akt signaling and promoting angiogenesis in human CRC. These findings not only provide the proteomic profiling of VECs in CRC, but also highlight some angiogenesis-related proteins, which may be the potential antiangiogenic targets and suggest new insights into the mechanism of angiogenesis in CRC.

## Materials and Methods

### Sample collection

Ten fresh-frozen samples of CRC and ten matched ANC tissues were taken from the Department of Surgery, Xiangya Hospital, Central South University, China, and used for iTRAQ labeling. The patients received neither radiotherapy nor chemotherapy before curative surgery and provided a written informed consent form for the study, which was approved by the local ethical committee. All specimens were obtained from surgical resection and stored at −80°C until further investigation.

An independent set of formalin-fixed paraffin-embedded tissue specimens, namely, 30 cases of nonneoplastic colonic mucosa (NCM), 30 cases of adenomatous colorectal polyps (AD), 30 cases of colorectal carcinoma in situ (CIS) and 50 cases of invasive colorectal carcinoma (ICC), were obtained from the Department of Pathology of the Xiangya Hospital at Central South University and used for immunohistochemical analysis.

### Laser capture microdissection

The frozen samples were prepared, stained and diagnosed by pathological examination and a rapid immunohistochemical staining technique was performed in less than 13 min with the DAKO rapid EnvisionTM–HRP system as described (Wu *et al*, 2010). To identify and visualize the VECs in human colorectal tissues for sequential LCM, the specific VEC marker CD31 was used for immuno-staining. After the blood vessels were apparent, the glass slides were replaced with LCM-specific glass slides. The stained areas that had the characteristic morphology of VECs were captured by LCM. LCM was performed with a Leica AS LMD system to purify the cells of interest from each type of tissue as previously described (Cheng *et al*, 2008; Li *et al*, 2012). Each captured cell population was over 95% homogeneous as determined by direct microscopic visualization.

### Trypsin digestion and labeling with iTRAQ reagents

To diminish the effects of biological variation on the proteomic results, equal amounts of proteins from ten cases of microdissected samples of CRC and ANC were pooled respectively. Two pooled protein samples were obtained for trypsin digestion and labeled with iTRAQ reagents as follows: CRC, labeled with iTRAQ 116, 117 and 118; ANC, labeled with iTRAQ 113, 114 and 115. The labeled digests were then mixed and dried.

### Off-line 2D LC-MS/MS

The mixed peptides were first separated on a strong cation exchange column into ten fractions according to the procedure described in our previous study (Mu *et al*, 2013). Each fraction was dried, dissolved in buffer C (0.1% formic acid, 5% acetonitrile), and analyzed on Q-Exactive systems (Thermo Scientific) in information-dependent mode. Briefly, peptides were separated on reverse-phase columns (EASY column, 10cm, ID 75μm, 3μm, C18-A2) with an Easy nLC system (Thermo Scientific). Peptides were separated by a linear gradient mobile phase A (5% acetonitrile, 0.1% formic acid) and mobile phase B (95% acetonitrile, 0.1% formic acid) from 5% to 40% of mobile phase B in 60 min at a flow rate of 300 nL/min. Survey scans were acquired from 300-1800 m/z with up to 20 precursors selected for MS/MS and dynamic exclusion for 60s.

### Data analysis

The software used for protein identification and quantitation was Proteome Discoverer 1.4 software. The peptide mass tolerance was set at ±20 ppm and the fragment tolerance mass was set at 0.1 Da. The data analysis parameters were set as follows: Sample type, Itraq 4 plex (peptide labeled); Cys alkylation, MMTS; Digestion, Trypsin; Instrument, Q-Exactive systems; Species, *Homo sapiens*; ID Focus, Biological modifications; Search Effort, Thorough; Max missed cleavages, 2; False discovery rate (FDR) Analysis, Yes; User Modified Parameter Files, No; Bias Correction, Auto; Background Correction, Yes. Identified proteins were grouped by the software to minimize redundancy. All peptides used for the calculation of protein ratios were unique to the given protein or proteins within the group, and peptides that were common to other isoforms or proteins of the same family were ignored.

The average iTRAQ ratios from the triple experiments were calculated for each protein. The confidence level of the altered expression of proteins was calculated by T-test as a P-value, which allows the results to be evaluated based on the confidence level of expression change.

### Bioinformatics analysis

The DEPs were first annotated by Gene Ontology (GO) using the PANTHER database (http://www.pantherdb.org/). Briefly, GO analysis was used to elucidate the genetic regulatory networks of interest by forming hierarchical categories according to the biological process, cellular component and molecular function aspects of the DEPs. Pathway analysis was performed to examine the significant pathways of the DEPs according to Metascape (http://metascape.org/gp/index.html#/main).

### Immunohistochemistry

Immunohistochemistry was performed according to the procedure described in our previous study (Zeng *et al*, 2012). Briefly, 4 μm thick tissue sections were deparaffinized, rehydrated, and treated with an antigen retrieval solution (10 mmol/L sodium citrate buffer, pH 6.0). The sections were incubated with anti-TNC (1:250; Abcam) or anti-CD34 (1:100; ZSGB-BIO) overnight at 4°C and then incubated with biotinylated secondary antibody followed by addition of avidin-biotin peroxidase. Finally, tissue sections were incubated with 3’,3’-diaminobenzidine until a brown color developed, and they were counterstained with Harris’ modified hematoxylin. In negative controls, primary antibodies were omitted. The evaluation of immunostaining was performed as previously described (Zeng *et al*, 2012). A score (ranging from 0-6) was obtained for each case. A combined staining score of ≤ 2 was considered to be weak staining (no/low expression), a score between 3 and 4 was considered to be moderate staining (expression), and a score between 5 and 6 was considered to be strong staining (high expression).

### Western blotting

For western blot analysis, the same amount (30ug) of whole cell lysate samples or enriched conditioned media samples were subjected to SDS-PAGE and subsequently transferred to PVDF membranes. Membranes were blocked in 5% milk for 2h prior to incubation with primary antibodies against TNC (1:1000; Santa Cruz), ITGB3 (1:1000; Proteintech), FAK Phospho-FAK Tyr397, Akt, Phospho-Akt Ser473 (1:1000; Cell Signaling Technology) overnight at 4°C. ACTB was used as a control for protein loading and was detected using a mouse anti-ACTB antibody (1:2000; Proteintech). Membranes were incubated with the corresponding secondary antibodies, including anti-rabbit and anti-mouse peroxidase (HRP)-linked IgG (1:1000; KPL) for 1h.

### Cell culture, cell transfection and conditioned media preparation

Human colon cancer cell lines SW620 and human umbilical vein endothelial cells (HUVECs) were maintained in RPMI-1640 or DMEM medium supplemented with 10% FBS in a humidified chamber with 5% CO_2_ at 37°C. SW620 cells and HUVECs were purchased from the Cell Bank of Type Culture Collection of the Chinese Academy of Sciences (Shanghai, China).

SW620 cells were cultured in 6-well plates until reaching 70% of confluency, the cells were transduced with lentiviral shRNA-TNC (GGAGTACTTTATCCGTGTATT) and lentiviral control (Cyagen Bioscience Inc., Guangzhou, China) at an MOI of 20 in the presence of 5 μg/ml polybrene for 24h and treated with 3 μg/ml puromycin for three days. The generated cell clones were tested for shRNA-TNC stable expression.

The untransfected (SW620), empty vector (SW620/Vector) and TNC shRNA plasmid-transfected SW620 (SW620/shTNC) cells were chosen for conditioned media preparation. Conditioned media were collected as previously described (Li *et al*, 2016). Briefly, approximately 3 × 10^6^ cells were grown to 70% confluency, and the medium was exchanged with serum-free RPMI-1640 medium. Conditioned media were collected 24h after the change of media and stored at −20°C for use. For subsequent experiments, the conditioned media were filtered using a 0.22 μm filter (Millipore) and concentrated using a Millipore centrifugal filter (3kDa). The protein concentration was determined by a standard Bradford protein assay (Thermo Scientific). After conditioned media were removed, the cell monolayer was washed, scraped and lysed in the presence of protease inhibitors. Protein concentration was determined using a Pierce BCA Protein Assay Kit (Thermo Scientific).

### HUVEC proliferation assay

The HUVECs were plated at 2×10^3^ cells per well in 96-well tissue culture plates and cultured in a 1:1 mixture of 10% FBS complete DMEM and conditioned media (total 200μl). The cells were cultured for 6 days. Every 24h, 20μl of CCK8 (5 mg/ml; Beyotime) was added to the wells, and cells were further incubated for 2h. The absorbance of each well was read with a Bio-Tek Instruments EL310 Microplate Autoreader at 450nm. The CCK8 assay was performed three times in triplicate.

### Wound healing assay

Cell migration was determined by scratch wound healing assay. Briefly, cells were grown in a 1:1 mixture of 10% FBS complete DMEM and conditioned media overnight to confluence in a 6-well plate. Monolayers of cells were wounded by dragging a pipette tip. Cells were washed to remove cellular debris and allowed to migrate for 12-24h. Images were taken at 0h and 12h after wounding under the inverted microscope.

### HUVEC migration assay

Migration activity was measured by Transwell assay (Corning, 3422). Approximately 5×10^4^ HUVECs were added to the upper chamber in 200μl of 1% FBS DMEM medium. The lower chamber contained 250μl of 10% FBS complete DMEM and 250μl of conditioned media. The plates were incubated for 24h at 37°C in 5% CO_2_. After incubation at 37 °C for 48h, cells were fixed with 4% paraformaldehyde and stained with 0.5% crystal violet. Each clone was plated in triplicate for each experiment, and each experiment was repeated at least three times.

### HUVEC angiogenesis assay

Matrigel (BD Biosciences, 354248) was melted at 4°C, added to 48-multiwell plates (Corning) at 100 μl/well, and then incubated at 37°C for 30 min. HUVECs (4×10^3^ cells) were resuspended in a 1:1 mixture of 250μl of 20% FBS complete DMEM and 250μl of conditioned media (total 500μl with 10% FBS). The cells were added to the wells, and after 7h incubation at 37°C, HUVEC tube formation was assessed by microscopy. Each well was imaged under a light microscope. The numbers of branches were calculated and quantified using MacBiophotonics ImageJ software (NIH).

### Statistical analysis

All statistical analyses were performed using the SPSS software package (version 13.0; SPSS, Inc.) and Prism 5.0 software (GraphPad). Data are shown as the mean ±SEM. The difference in TNC protein expression between the two different kinds of tissue (CRC vs. ANC) was analyzed using a linear trend chi-square test. The relationship between TNC expression and the clinicopathological characteristics in patients with CRC was analyzed using the Mann-Whitney U test and linear trend chi-square test. For *in vitro* assays, the data are reported as biological replicates, with technical replicates indicated in the figure legends. Student’s t-tests (unpaired, two-tailed) were performed to determine whether a difference between two values was statistically significant, with P < 0.05 considered significant. *In vitro* assays were performed in triplicate unless otherwise stated. *p* values < 0.05 were considered statistically significant (\**p* < 0.05; \*\**p* < 0.01; \*\*\**p* < 0.001).

## Acknowledgements

We would like to thank Prof. Hongwei Lv and Prof. Hong Xiang of Third Xiangya Hospital for the introduction to the HUVECs angiogenesis assay.

